# Error-Controlled Discovery of Protein Interactions via Conformal Calibration

**DOI:** 10.64898/2026.11.03.764142

**Authors:** Yifan Yang, Dehan Li, Daoyun Ding, Lingfeng Shi, Han Zhou, Yang Young Lu

**Author notes:** These authors contributed equally to this work (Yifan Yang, Dehan Li, Daoyun Ding).

## Abstract

Protein language models (PLMs) are powerful predictors of protein interactions, but strong benchmark performance does not ensure reliable discoveries. We introduce PPI-Caliper, a model-agnostic framework that adapts conformal calibration to select interactions at user-specified false discovery rate (FDR) levels using held-out labels without retraining the predictor. Across five species, two PLM-based predictors, and two human training-data versions for each predictor, PPI-Caliper keeps empirical FDR at or below the target level under in-distribution calibration. Under distribution shift, transferred thresholds tend to become conservative when predictive performance improves, but can fail to control FDR when performance deteriorates. PPI-Caliper connects pre-trained interaction scores to explicit error targets and demonstrates why reliable biological discovery requires evaluating calibration alongside predictive accuracy.

## 1 Introduction

Protein–protein interactions (PPIs) organize proteins into complexes and networks that coordinate cellular functions [1]. Uncovering these interactions is therefore essential for decoding disease mechanisms and guiding drug discovery [2]. Because experimental interaction identification is costly, computational methods provide a scalable way to prioritize candidate interactions for validation, particularly when interaction maps remain incomplete. Protein language models (PLMs) have emerged as a powerful solution for PPI prediction by learning structural and functional representations from large collections of protein sequences [3, 4]. These representations can then be combined, or sometimes jointly modeled and refined, to predict whether two proteins interact [5, 6], making it increasingly practical to rank candidate interactions across proteins and species.

However, strong benchmark performance does not ensure that individual predictions are trustworthy enough to guide experimental follow-up. Benchmark metrics such as AUROC and AUPRC summarize overall predictive performance, but do not indicate how many interactions selected at a given threshold are false. This limitation is especially important when non-interacting protein pairs dominate the candidate set, where even a low false-positive rate can yield many incorrect predictions relative to true interactions [7]. The key discovery question is therefore not simply whether interacting protein pairs receive higher scores, but how many predicted interactions can be reported reliably. False discovery rate (FDR) control addresses this question by limiting the expected fraction of false positives among selected interactions [8, 9]. For example, an FDR threshold of 1% means that, on average, no more than 1% of reported interactions are expected to be false, providing an interpretable basis for experimental follow-up rather than relying on benchmark performance or an arbitrary prediction score cutoff.

Estimating FDR requires a reference for the scores expected from false predictions. A standard way is to model a null distribution and use it to derive p-values for multiple-testing correction [8]. Other settings, including proteomics database search [10, 11] and knockoff inference [12–15] instead construct synthetic negative controls that provide an empirical reference for false discoveries. However, applying these strategies to PPI prediction is challenging for two reasons. First, each candidate protein pair receives a single interaction score, without a natural null distribution against which that score can be compared. Second, synthetic negatives must realistically resemble natural non-interacting proteins, since PLMs can readily distinguish shuffled or reversed sequences and thereby underestimate false discoveries [16]. Conformal calibration provides an alternative by using held-out labeled data to directly select prediction thresholds that control false discoveries [17, 18]. Assuming that the calibration protein pairs are representative of those encountered at deployment, this avoids explicitly modeling a null distribution or constructing synthetic decoys. A similar strategy has already been successfully applied to control false positives in enzyme function annotation [19].

Here, we introduce PPI-Caliper, a model-agnostic framework that adapts this calibration strategy to obtain FDR-controlled interaction sets from existing PPI prediction models (Figure 1). PPI-Caliper first transforms prediction scores into calibration percentiles, making thresholds comparable across datasets even when their score distributions shift. It then resamples held-out labeled protein pairs to estimate the false discovery proportion (FDP) at candidate thresholds. Using a “Learn then Test” procedure [20], PPI-Caliper selects the least restrictive threshold that satisfies the target FDR without retraining the PPI prediction models. We evaluate PPI-Caliper under both in-distribution calibration, where calibration and test data come from the same species, and distribution-shifted transfer, where a threshold calibrated on one species is applied to another. Across datasets from five species and two widely used PLM-based PPI prediction models trained on human data, PPI-Caliper consistently controls empirical FDR at the target level in the in-distribution setting. Under distribution shift, however, transferring the calibrated threshold does not always preserve FDR control. When the threshold is transferred to a dataset where the prediction model performs better, it tends to be overly conservative, yielding fewer discoveries while keeping the empirical FDR below the target. In contrast, when prediction model performance degrades on the new dataset, the transferred threshold can underestimate the error rate, causing the empirical FDR to exceed the target. We further show that shifts in predictive performance and score distributions provide useful diagnostic signals for whether a calibrated threshold is likely to transfer reliably, offering a practical way to screen for potential failures before deployment. Together, these results establish PPI-Caliper as a practical approach for post-hoc FDR control in PPI prediction and provide guidance for applying calibration under distribution shift.

**Fig. 1:**
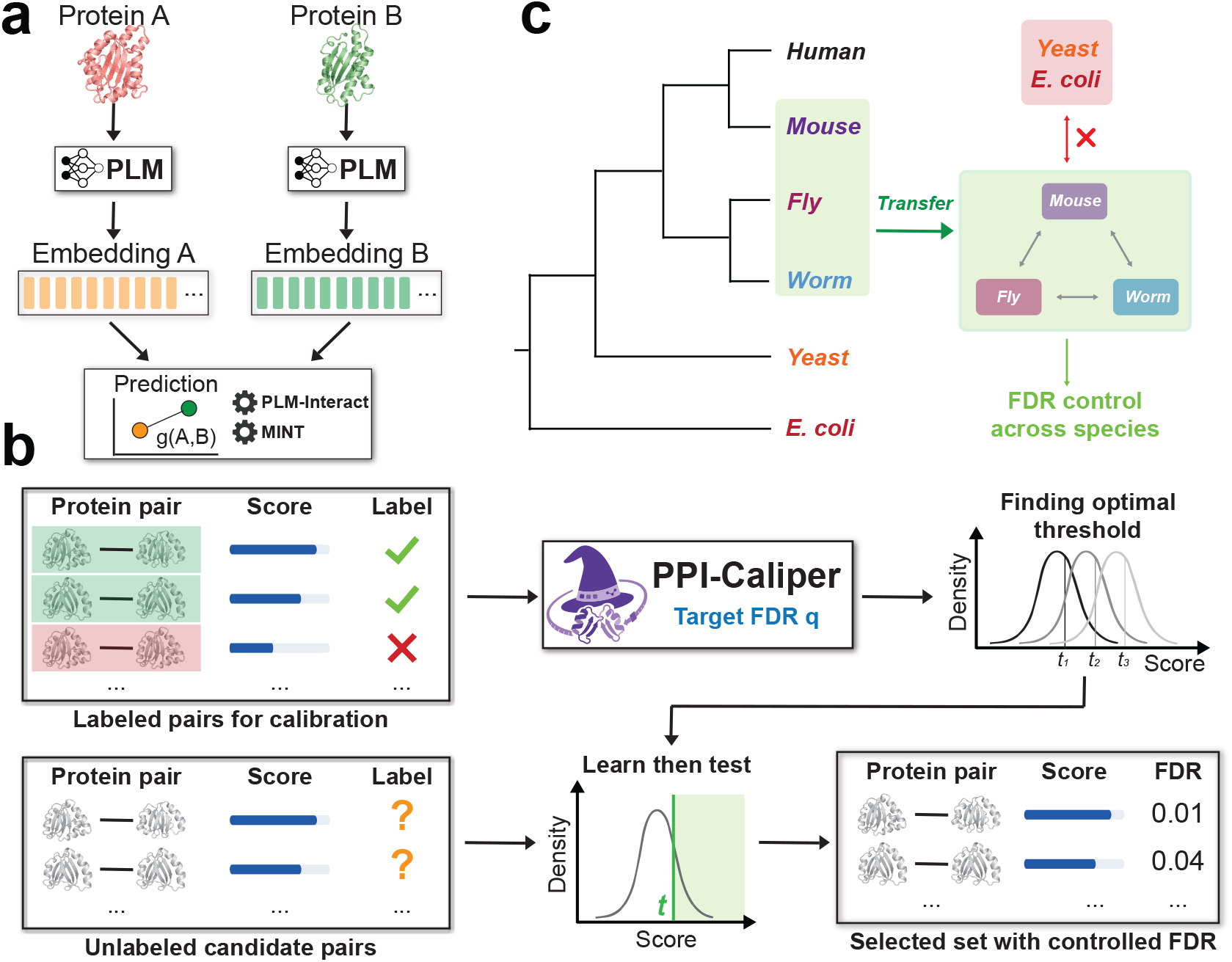
PPI-Caliper calibrates PPI scores for error-controlled discovery. **(a)** A PLM-based prediction model scores each protein pair and ranks candidate interactions. **(b)** PPI-Caliper estimates FDP from held-out labeled protein pairs and uses Learn then Test to select a threshold *t* for a target FDR *q*, yielding FDR-controlled interaction sets. **(c)** Under distribution shift, thresholds transfer more reliably among mouse, fly, and worm than between these species and yeast or *E. coli*.

## 2. Related work

PPI-Caliper works alongside existing computational PPI prediction methods by calibrating their output scores. Earlier computational methods predict interactions from sequence, evolutionary, or structural features, typically by training classifiers to distinguish interacting from non-interacting protein pairs [21, 22]. PLM-based methods instead use representations learned from large protein sequence collections to predict interactions directly. D-SCRIPT combines pre-trained protein representations with a structure-aware contact module [23], TUnA builds on ESM-2 [4] to model predictive uncertainty [24], PLM-Interact fine-tunes ESM-2 to jointly encode protein pairs [5], and MINT captures interactions through cross-chain attention [6]. These methods improve PPI prediction performance and uncertainty estimation, while our work addresses the complementary question of which predictions can be reported reliably at a specified FDR level.

PPI-Caliper uses conformal calibration to improve the reliability of protein discovery. Related work has applied similar ideas to protein search, where Boger et al. [18] calibrate score thresholds to control FDR without retraining the underlying model, and to enzyme annotation, where CPEC uses conformal calibration to limit false functional assignments [19]. For PPI screening, Predictomes estimates FDR from curated reference pairs [25], while FlashPPI chooses a contact-score threshold to achieve a target precision on an importance-weighted benchmark [26]. Conformal methods have also been developed for link prediction, where FDR is controlled over missing edges in partially observed networks under assumptions about how edges are observed [27, 28]. PPI-Caliper addresses a different setting. It provides post-hoc FDR calibration for pairwise PPI predictions made directly from protein sequences, without requiring a network observation model, and explicitly studies whether calibrated error control remains reliable when transferred across shifted species datasets.

Lastly, PPI-Caliper builds on a broader line of work that combines flexible machine learning with explicit control of false discoveries. Model-X knockoffs provide a general framework for identifying informative variables with FDR control while allowing the predictive model itself to be highly flexible [12, 13]. DeepPINK extends this idea to deep neural networks by jointly modeling original and knockoff features for error-controlled variable selection [14]. Diamond further uses knockoff controls to identify non-additive feature interactions while separating joint effects from individual feature contributions [15]. Together, these methods show how statistical error control can be integrated with modern machine learning to make discoveries more reliable. Their primary goal is to determine which input features or feature combinations are associated with an outcome. Our setting is complementary and focuses on deciding which predictions from an existing PPI model can be reported while controlling the FDR.

## 3. Approach

### 3.1 Problem setup

We consider selecting reliable interaction predictions from an existing PPI prediction model. Let *x*_*i*_ = (*a*_*i*_, *b*_*i*_) denote a protein pair and *y*_*i*_ *∈ {*0, 1*}* its interaction label. The PPI prediction model *g* assigns each protein pair a score *g*(*x*_*i*_), with larger values indicating stronger evidence of interaction. We use a labeled calibration set *C*, held out from prediction model training, to choose a score threshold and apply it to a separate candidate set *T* for interaction discovery. Such calibration data can be drawn from existing PPI benchmarks, which contain experimentally supported interactions and sampled negative ones [5, 6]. Calibration does not change the prediction model, and labels in *T*, when available, are used only for evaluation.

Let *u*_*i*_ denote the normalized prediction score defined below. For the candidate protein pairs in *T*, a threshold *t* selects *R*(*t*) = {*i* | *u*_*i*_ *≥ t}* . The false discovery proportion (FDP) is the fraction of selected protein pairs labeled as non-interacting, and the false discovery rate (FDR) is its expectation:

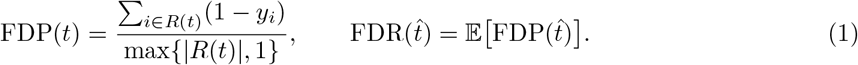

We define the FDP to be zero when no protein pairs are selected. Given a user-specified target *q ∈* [0, 1], our goal is to choose a threshold 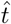that selects as many candidate interactions as possible while satisfying 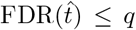. The expectation accounts for variation in both the data and the calibration procedure, providing a set-level error criterion for the reported interactions.

### 3.2 Score-to-quantile normalization

Raw prediction scores can vary substantially across datasets, making a single numerical threshold difficult to compare or transfer. We therefore convert each score to its percentile relative to the calibration-score distribution. Specifically, we compute empirical score quantiles at 1,000 evenly spaced levels using only scores from *C* and use the resulting score–quantile pairs to construct a monotone mapping *ϕ*. Each prediction score is then transformed as:

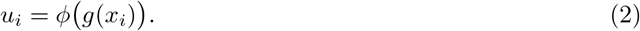

After fitting, the mapping is applied to candidate protein pairs in *T*. Because the transformation is monotone, it preserves score ordering and changes only the scale on which thresholds are expressed. We then use the calibration labels to determine which thresholds support controlled selection error.

### 3.3 Selection error estimation

Having placed prediction scores on a shared scale, we next use the labeled calibration set to estimate the false discovery proportion associated with each candidate threshold. We construct *K* bootstrap batches *B*_1_, …, *B* _*K*_, by sampling *m* labeled protein pairs from *C* with replacement for each batch. Each batch serves as a small candidate set on which we evaluate the same threshold-based selection rule. We consider a grid *G* of thresholds spanning the normalized scores observed in these batches. For each threshold *t ∈ G*, we select pairs with *u*_*i*_ *≥ t* in each batch and compute the resulting false discovery proportion FDP_*k*_(*t*) using Eq. 1. These batches provide repeated measurements of selection error at each threshold. We summarize them by their mean and standard deviation:

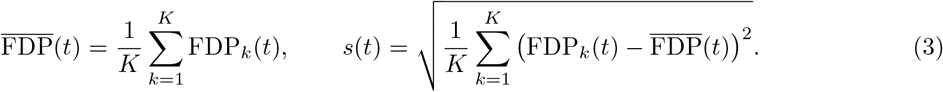

The mean estimates the expected false discovery proportion for a batch of *m* pairs, while the standard deviation describes variation across resampled batches. Together, they allow us to assess a threshold using both its average selection error and the variability of that error.

### 3.4 Threshold selection

Ideally, if higher score thresholds always reduced FDP, we could simply raise the threshold until the target error level was met. However, stricter thresholds can also remove true interactions, so FDP is not guaranteed to decrease. We therefore use Learn then Test (LTT), which evaluates thresholds without assuming that higher thresholds always yield lower FDP [20]. For each threshold, the null hypothesis is that its expected batch FDP exceeds the target *q*. A threshold passes the test when the calibration data provide sufficient evidence against this hypothesis. Following the implementation used in conformal protein retrieval [18], we compute the normal-approximation p-value:

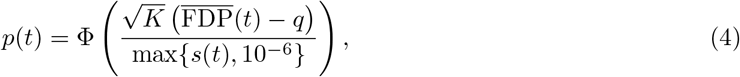

where Φ is the standard normal cumulative distribution function and the denominator safeguard prevents division by zero. The test compares how far the mean batch FDP lies below *q* with its estimated variability. A threshold passes when *p*(*t*) *≤ δ*, where *δ* is the testing level rather than the target FDR. To select among thresholds, we use a fixed-sequence scan from the largest threshold downward. We begin with the highest-scoring protein pairs and progressively admit additional protein pairs while the corresponding thresholds pass the test. The scan stops at the first failure, and we select the last passing threshold. Equivalently, when at least one threshold is accepted,

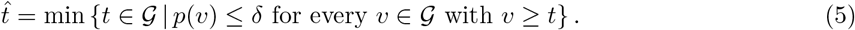

This rule selects the least restrictive threshold reached by the scan, rather than continuing to search for isolated passing thresholds after a failure. We then apply 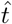 to *T* and report the selected interaction set 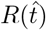. The batch statistics are computed once and can be reused for different target levels *q*. Under valid risk tests and an appropriate testing sequence, LTT bounds by *δ* the probability of accepting any threshold whose expected batch FDP exceeds *q* [20]. We use bootstrap batches drawn from the same finite calibration set and a normal approximation for their mean FDP. These approximations do not by themselves establish a finite-sample population FDR guarantee. Moreover, expected batch FDP can differ from FDR on the complete candidate set, as discussed in Appendix B. We therefore assess the resulting FDR empirically on held-out labeled pairs.

## 4 Results

### 4.1 Experimental setup

#### 4.1.1 Evaluation design

We evaluated PPI-Caliper in two complementary settings. In the in-distribution setting, calibration and test data come from the same species, allowing us to assess whether the selected interactions achieve the target FDR under matched distributions. In the distribution-shifted setting, we transfer a threshold calibrated on one species to another and test whether FDR control remains valid as the data distribution changes. Each species-specific dataset defines a distinct evaluation domain. Importantly, these settings differ only in the relationship between the calibration and test distributions, not in the data used to train the underlying PPI predictor.

#### 4.1.2 Evaluation datasets and models

We evaluated PPI-Caliper on five PPI datasets from mouse, fly, worm, yeast, and *E. coli*, using PLM-Interact [5] and MINT [6] as the underlying prediction models. For PLM-Interact, we used the released humanV11 checkpoint trained on a human benchmark derived from STRING V11 [29], and the humanV12 checkpoint trained on a separate human PPI dataset constructed from STRING V12 [30]. We fine-tuned MINT on the same two human datasets to obtain matched checkpoints. Together, these four model and training-data combinations allow us to assess whether calibration and transfer behavior is consistent across prediction architectures and supervised training datasets. All prediction models remain unchanged during calibration and evaluation.

#### 4.1.3 Evaluation procedure

For in-distribution evaluation, we repeated the experiment 100 times for each dataset and model check-point. For each repetition, we randomly split the labeled pairs equally, stratified by interaction label, into a calibration set *C*_*s*_ and a held-out test set *T*_*s*_. For distribution-shifted evaluation, we instead used the source calibration set *C*_*s*_ to determine the score-to-quantile mapping and threshold, and applied them to the held-out test set *T*_*t*_ of a different target species. Thus, the first setting evaluates calibration on held-out data from the same domain, while the second evaluates transfer to a different domain. In both settings, labels in test sets were used only for evaluation and never for threshold selection. We fit the score-to-quantile mapping *ϕ* on *C* using 1,000 equally spaced quantile levels, then drew *K* = 10 boot-strap calibration batches, each containing *m* pairs, where *m* was set to 10% of the source dataset before splitting. We considered target FDR levels *q ∈ {*0.1, 0.2, …, 0.9 } and an equally spaced threshold grid *G* spanning the normalized scores in the bootstrap batches. For each target level, we selected 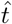 using the Learn then Test procedure in Section 3 and applied it to *T* to obtain the selected interaction set 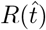.

#### 4.1.4 Evaluation metrics

We computed the FDP of 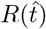 using Eq. 1 and reported its mean across repetitions as the empirical FDR. Power is defined as the fraction of true interactions in *T* that are selected. We also reported classification accuracy, which varies with the calibrated threshold by treating selected pairs as interactions and all remaining pairs as non-interactions. To characterize the underlying predictive performance independently of threshold selection, we reported AUPRC for each model and dataset, following the evaluation used for PLM-Interact. We summarized FDR, power, and accuracy by their means and 95% confidence intervals over 100 repetitions.

### 4.2 FDR control under in-distribution calibration

We first evaluated whether PPI-Caliper achieves the target FDR when calibration and test data come from the same distribution. For PLM-Interact humanV11, empirical FDR remains close to or slightly below the target across all five datasets, while power increases steadily as the allowed FDR level increases (Figure 2). This shows that PPI-Caliper allows users to recover more interactions by allowing a higher FDR, while keeping the observed error rate close to the specified target.

**Fig. 2:**
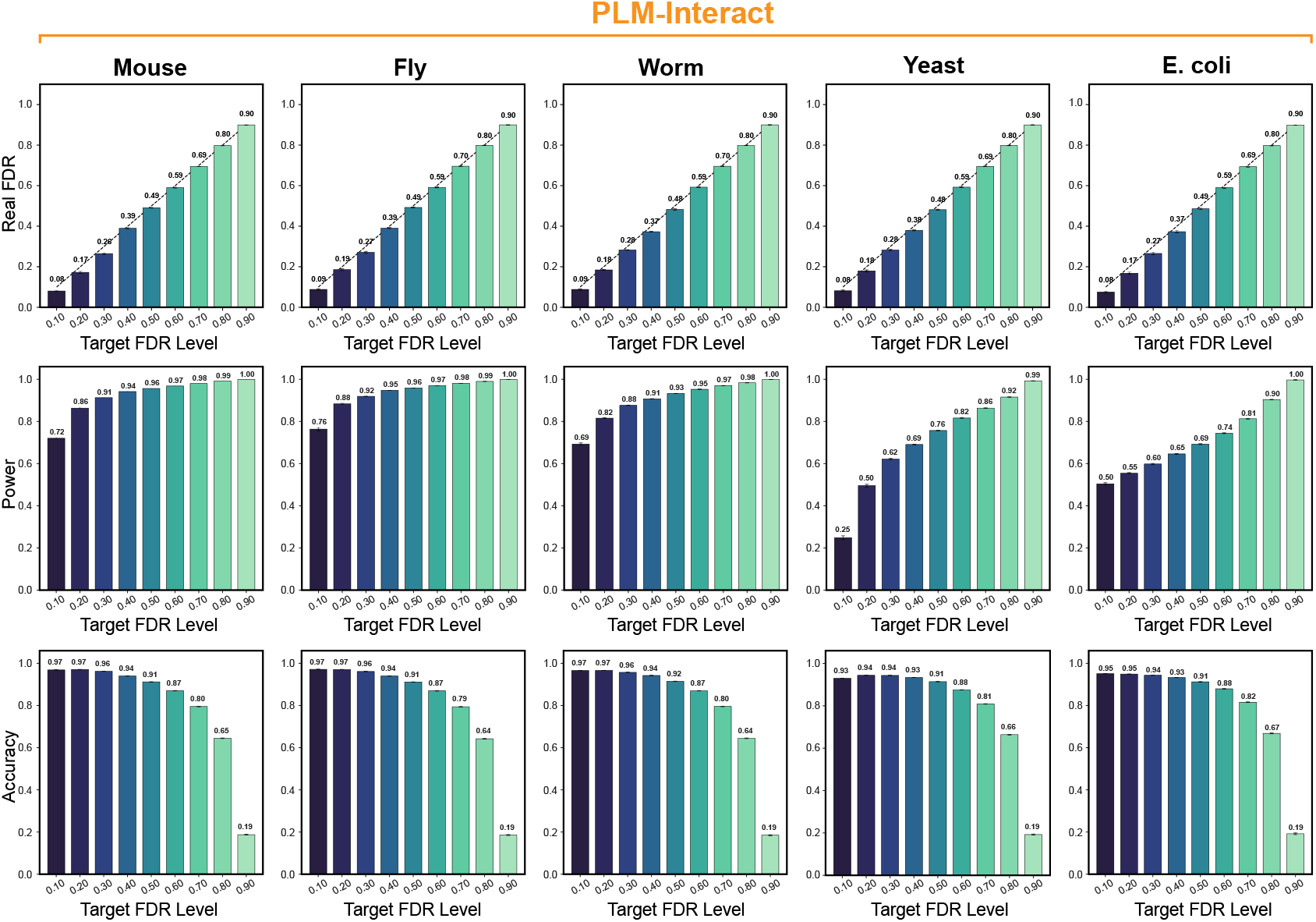
In-distribution calibration controls FDR across datasets with varying discovery power. Results are shown for PLM-Interact humanV11 across five datasets. Rows report empirical FDR, power, and accuracy over target FDR levels *q*. Bars show means over 100 random 50%/50% calibration and test splits with 95% confidence intervals. The dashed line marks empirical FDR equal to the target. Power is the fraction of true interactions in the test set that are selected.

Although FDR is controlled consistently, discovery power varies substantially across datasets. At *q* = 0.1, power ranges from 0.25 on yeast to 0.76 on fly, with stronger ranking performance and higher power on mouse, fly, and worm than on yeast and *E. coli*. The same pattern holds for PLM-Interact humanV12 and both matched MINT checkpoints (Figures A1, A2, and A3). Across these model and training-data settings, empirical FDR remains close to the target FDR level even when power differs substantially. Together, these results show that PPI-Caliper can control FDR across datasets and model variants, while the number of true interactions recovered still depends on how well the underlying PPI prediction model performs.

Classification accuracy is less informative about these changes in discovery performance. For PLM-Interact humanV11, accuracy remains between 0.91 and 0.97 as *q* increases from 0.1 to 0.5, even though empirical FDR rises from approximately 0.08 to 0.49. This occurs because the benchmark contains ten sampled negative pairs for every positive pair, so correctly classified negatives dominate overall accuracy and can hide changes in the reliability of the selected interactions. Accuracy drops substantially only when the target FDR reaches *q* = 0.9, falling to approximately 0.19. Empirical FDR and power therefore provide a more informative view by directly measuring how reliable the selected interactions are and how many true interactions are recovered.

### 4.3 FDR control under distribution-shifted transfer

#### 4.3.1 Transfer reliability depends on direction

Successful in-distribution calibration does not guarantee that the same threshold will control FDR after the data distribution changes. We therefore transferred each source-calibrated threshold 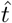 to a different target dataset without using target labels for recalibration. Figure 3 summarizes these experiments for PLM-Interact humanV11, with diagonal panels showing in-distribution calibration and off-diagonal panels showing threshold transfer across datasets.

**Fig. 3:**
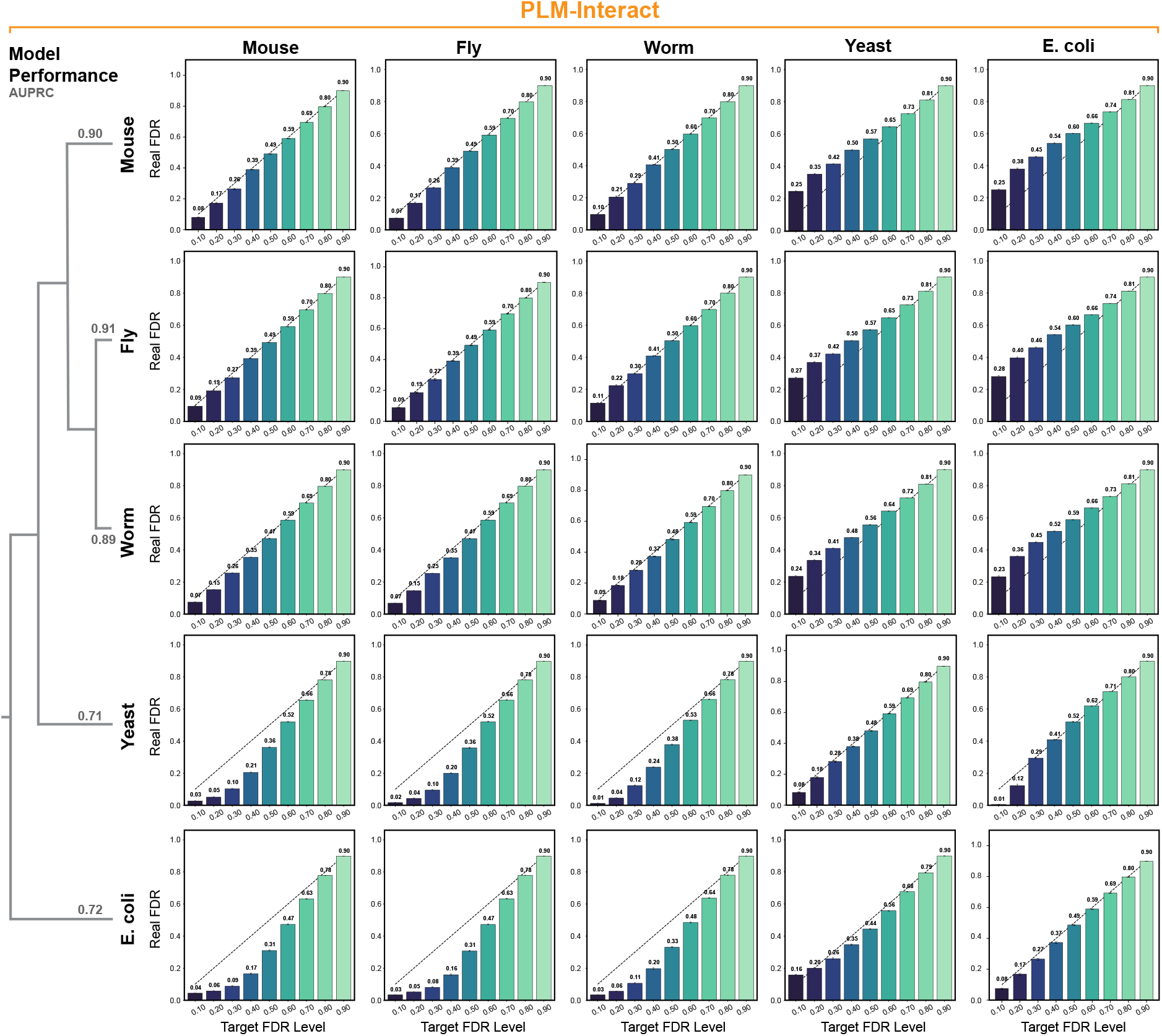
FDR control under distribution shift depends on transfer direction. Results are shown for PLM-Interact humanV11. Rows indicate calibration datasets and columns indicate test datasets, with diagonal panels showing in-distribution evaluation. Bars show mean FDP over 100 repetitions with 95% confidence intervals, and the dashed line marks the target FDR. AUPRC values summarize prediction performance on each dataset.

Transfer behavior follows a clear pattern across datasets and depends strongly on how prediction performance changes between the source and target domains. Thresholds transferred among mouse, fly, and worm generally keep empirical FDR close to the target FDR level. When a threshold is calibrated on a lower-performing dataset such as *E. coli* and transferred to a higher-performing dataset, empirical FDR usually remains below the target, although the threshold may select fewer interactions than necessary. In the reverse direction, transferring a threshold from a higher-performing dataset to a lower-performing one can underestimate the error rate and cause empirical FDR to exceed the target. Thus, transfer reliability depends on the direction of the performance shift. A threshold may remain safe but become overly restrictive when performance improves, whereas FDR control can fail when performance deteriorates. Reliable calibration in the source domain therefore does not guarantee FDR control after transfer.

#### 4.3.2 Predictive performance provides a diagnostic for calibration transfer

We next examine whether changes in predictive performance can help identify when calibration is likely to transfer reliably. PLM-Interact achieves substantially higher AUPRC on mouse, fly, and worm than on yeast and *E. coli* (Figure 3). Thresholds transfer more reliably between datasets with similar predictive performance, while larger performance gaps are associated with larger deviations from the target FDR. The direction of the change also matters. Improved performance after transfer tends to produce more conservative selection, whereas degraded performance increases the risk that empirical FDR exceeds the target.

This relationship is consistent across model variants. PLM-Interact humanV11 and humanV12 show similar performance differences across datasets and corresponding transfer behavior (Figures 3 and A4), and both MINT checkpoints show the same overall trend (Figures A5 and A6). Changes in score distributions provide an additional signal. Although the monotone mapping *ϕ* places scores on a common percentile scale, true and false interactions may still occupy different score regions after a distribution shift. The same normalized threshold can therefore select different proportions of true and false interactions across datasets.

Together, these results suggest a practical way to screen calibration transfer before deployment. A calibrated threshold is more likely to remain reliable when source and target datasets have similar predictive performance and score distributions, while larger shifts indicate a greater risk of losing FDR control. Predictive performance and score-distribution changes can therefore provide useful signals for deciding whether recalibration may be needed in a new domain.

## 5. Discussion and conclusion

PPI-Caliper addresses the gap between accurate PPI prediction and reliable interaction selection for bio-logical discovery. It calibrates existing PPI predictors using held-out labeled pairs without modifying the underlying model. Across five evaluation datasets, two predictors, and two human training-data settings, PPI-Caliper keeps empirical FDR at or below the target level under in-distribution calibration, although discovery power varies substantially. These results highlight the complementary roles of prediction and calibration. Better predictors recover more true interactions, while calibration connects their selection to an explicit error target. Evaluating FDR together with power provides a more informative view of discovery performance than benchmark accuracy alone.

The transfer experiments show that a reliable threshold in one domain may not remain reliable after distribution shift. When predictive performance improves after transfer, the threshold tends to become conservative. When performance deteriorates, the transferred threshold can underestimate error and exceed the target FDR. Changes in predictive performance and score distributions therefore provide useful signals of potential calibration mismatch, although neither alone guarantees reliable transfer. Our conclusions are also limited by the current calibration procedure and benchmark labels. Bootstrap estimates of batch-level FDP do not provide a finite-sample guarantee for the full candidate set, and sampled negatives may contain unobserved interactions. The current results support practical calibration on the evaluated distributions rather than guarantees in new biological screens.

Future work should strengthen both the statistical guarantees and robustness of calibration under distribution shift. Direct conformal selection could target FDR over the full candidate set without the current batch-level approximation [31], although its assumptions should be examined for protein pairs that share proteins and for limited calibration data. Under distribution shift, weighted conformal methods could adjust calibration when protein-pair features change while the conditional relationship with interaction labels remains stable [32]. Broader evaluation should include additional prediction models such as D-SCRIPT [23] and TUnA [24], smaller calibration sets, and splits that hold out entire proteins or sequence families to reduce known benchmark biases [33].

## Data Availability

The protein–protein interaction datasets and supporting data for the analyses and figures reported in this study are publicly available in the PPI-Caliper GitHub repository at https://github.com/batmen-lab/PPI-Caliper/tree/main/data. Instructions for downloading and preparing these data are provided in the repository documentation.

## Code Availability

Source code for PPI-Caliper and all reported analyses is publicly available in the GitHub repository at https://github.com/batmen-lab/PPI-Caliper. Setup and reproduction instructions are provided in the repository documentation.

## A Experimental details

### A.1 Datasets

We evaluate on the five-species cross-species PPI benchmark of Sledzieski et al. [23], distributed with PLM-Interact [5]. The benchmark contains mouse, fly, worm, yeast, and *E. coli*. Positive pairs correspond to experimentally supported physical interactions from STRING. To reduce overlap with the human training data, proteins highly similar to human training proteins were removed by clustering at 40% sequence identity with CD-HIT. Negative examples are randomly sampled protein pairs treated as non-interacting by the benchmark, at a ratio of ten negatives per positive. Table 1 summarizes the resulting dataset sizes.

**Table 1:**
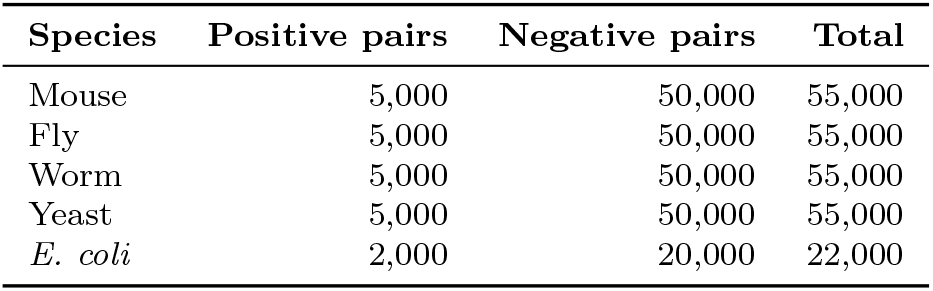
Per-species benchmark sizes.

| **Species** | **Positive pairs** | **Negative pairs** | **Total** |
|---|---|---|---|
| Mouse | 5,000 | 50,000 | 55,000 |
| Fly | 5,000 | 50,000 | 55,000 |
| Worm | 5,000 | 50,000 | 55,000 |
| Yeast | 5,000 | 50,000 | 55,000 |
| *E. coli* | 2,000 | 20,000 | 22,000 |

### A.2 Prediction models

We evaluate two PLM-based PPI predictors, PLM-Interact [5] and MINT [6]. For PLM-Interact, we use the released humanV11 and humanV12 checkpoints. The humanV11 checkpoint is trained on the D-SCRIPT human benchmark derived from STRING V11, whereas humanV12 is trained on the human PPI dataset constructed by the PLM-Interact authors from STRING V12. For a matched comparison, we fine-tune MINT separately on the corresponding V11- and V12-based human PPI datasets. All calibration procedures operate on the output scores of these fixed predictors and do not update their parameters.

### A.3 Within-species calibration

For each species and model checkpoint, we perform 100 independent trials. In each trial, the labeled dataset *D*_*s*_ is randomly divided 50/50, stratified by interaction label, into equal-sized calibration and held-out test sets, denoted by *C*_*s*_ and *T*_*s*_, respectively. The calibration set is used for score normalization and threshold selection, while labels in *T*_*s*_ are used only to evaluate the selected interaction set. For each trial, we fit the score-to-quantile mapping *ϕ*_*s*_ using only prediction scores from*C*_*s*_. We compute score values at 1,000 evenly spaced quantile levels in [0, 1] and use them to construct the monotone mapping described in Section 3.2. The same mapping is then applied to scores in both *C*_*s*_ and *T*_*s*_, and no test labels or test scores are used to construct the mapping. We draw *K* = 10 bootstrap calibration batches from *C*_*s*_ uniformly with replacement, without stratifying by label. Each batch contains round(|*D*_*s*_ |*/*10) protein pairs, that is, 5,500 pairs for mouse, fly, worm, and yeast, and 2,200 pairs for *E. coli*. For each candidate threshold, we compute the FDP independently within each batch, defining it as zero when no pair is selected, and summarize the resulting *K* values by their mean and standard deviation. The standard deviation is floored at 10^*™*6^, and the p-values in Eq. equation 4 use a sample size of *K*. Thresholds are evaluated over a grid of *N* = 100 equally spaced values between the smallest and largest normalized scores observed across the *K* bootstrap batches. We set the LTT testing level to *δ* = 0.5, following the default of the get thresh FDR routine of Boger et al. [18]. Under this choice, the p-value of a grid threshold is at most *δ* exactly when its mean batch FDP does not exceed the target level *q*, so the selected threshold is the smallest grid value above which the bootstrap mean FDP stays at or below *q*. For each target FDR level *q ∈ {*0.1, 0.2, …, 0.9*}*, we compute the normal-approximation p-values in Eq. equation 4 and apply the fixed-sequence procedure in Eq. equation 5. The selected threshold is then applied to the held-out test set *T*_*s*_. Algorithm 1 summarizes the complete calibration procedure.

### A.4 Cross-species transfer

To evaluate robustness under distribution shift, we transfer calibrated thresholds between species, considering all 20 ordered source–target pairs. In each of the 100 trials, every species *s* is split equally, stratified by interaction label, into a calibration set *C*_*s*_ and a held-out test set *T*_*s*_, and the same split is shared by the within-species and cross-species experiments. For a source species *s*, the score-to-quantile mapping *ϕ*_*s*_ and the threshold are determined from *C*_*s*_, exactly as in the corresponding within-species experiment. The source-fitted mapping *ϕ*_*s*_ is then applied to the prediction scores of the held-out target set *T*_*t*_, and the source-calibrated threshold is applied to the resulting normalized scores. Empirical FDR is evaluated on *T*_*t*_. Thus, for a fixed trial, all transfers from the same source species share the same score-to-quantile mapping and calibrated threshold.

### A.5 Evaluation metrics

For within-species experiments, we report empirical FDR, power, and classification accuracy on the heldout test set, averaged over the 100 random calibration–test splits with 95% confidence intervals. Power is defined as the fraction of labeled positive interactions recovered by the selected interaction set. For crossspecies experiments, we report empirical FDR as a function of the target FDR level for every ordered source–target species pair. We additionally report area under the precision–recall curve (AUPRC) to characterize the underlying predictive performance of each model on each species.

## B Relationship between batch-level FDP and candidate-set FDR

Our calibration procedure estimates FDP on bootstrap batches of size *m*, whereas the selected threshold is ultimately applied to a larger candidate set. These quantities differ because the FDP of an empty selected set is defined as zero.

Fix a threshold *t*. Let

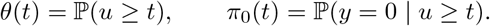

For a batch of *m* independently sampled pairs, the number of selected pairs satisfies *S*_*m*_(*t*) *∼*

Binomial(*m, θ*(*t*)). Conditional on *S*_*m*_(*t*) *>* 0, the expected FDP is *π*_0_(*t*). Therefore,

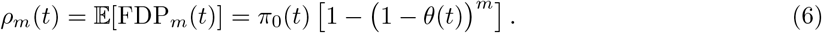

Similarly, for a candidate set of size *n* drawn from the same distribution,

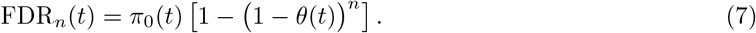

Thus, when *n ≥ m*, the candidate-set FDR is at least as large as the batch-level expected FDP, with the difference arising only from the probability of selecting no pairs. This difference becomes negligible when *mθ*(*t*) is sufficiently large.

Under cross-species transfer, the selection rate *θ*(*t*) and conditional error rate *π*_0_(*t*) can both change between the calibration and target distributions. Consequently, the transferred threshold need not preserve FDR control even when the empty-selection effect is negligible. Our implementation additionally uses overlapping bootstrap batches from a finite calibration set, so we assess candidate-set FDR empirically on held-out data.

## C PPI-Caliper algorithm

Algorithm 1 summarizes the complete calibration and threshold-selection procedure described in Section 3; the batch-level FDP statistics are computed once and reused across target FDR levels.

### Algorithm 1

PPI-Caliper calibration and threshold selection

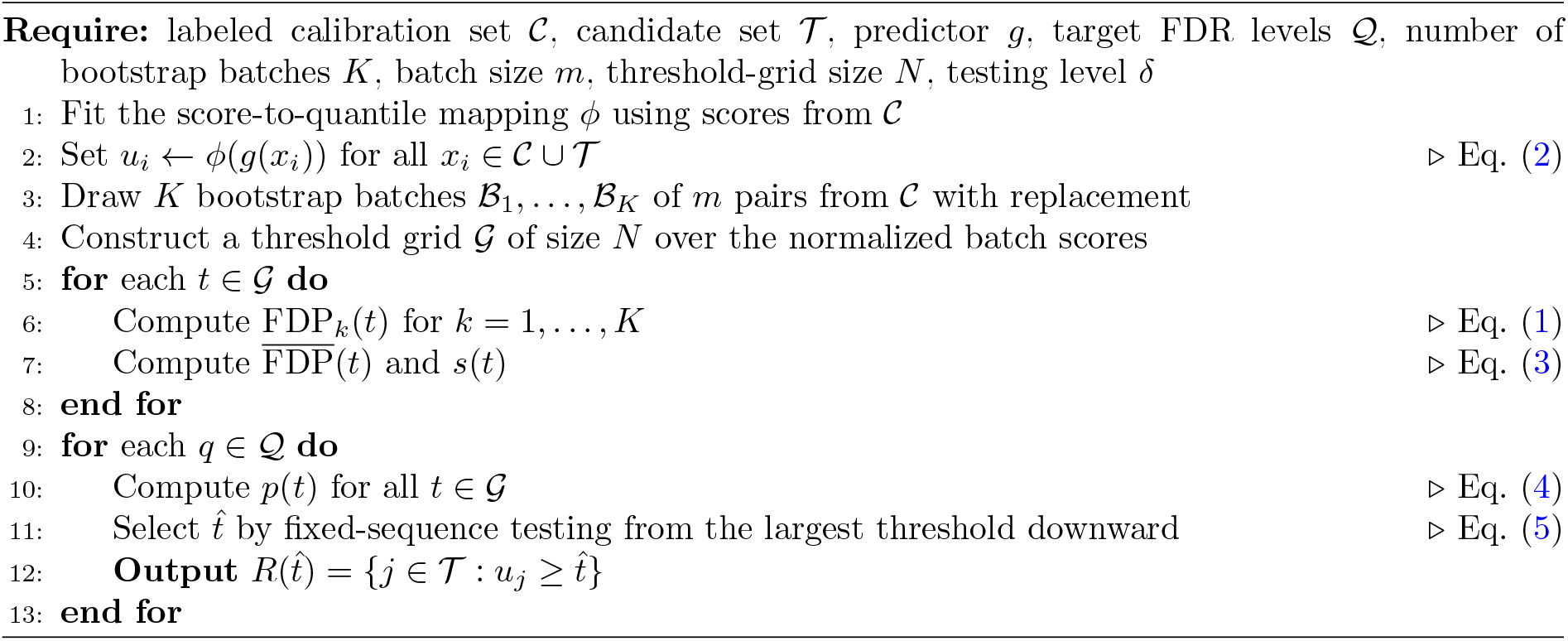

## D Additional results

This appendix repeats the experiments of Section 4 for the humanV12 checkpoint of PLM-Interact and for MINT [6], which we fine-tuned separately on the STRING V11 and V12 human PPI datasets. Figures A1–A3 show within-species control, and Figures A4–A6 show cross-species transfer. All panels follow the layout of Figures 2 and 3 and report the mean over random splits with 95% confidence intervals. The findings of the main text hold across these additional model and training-data combinations: the realized FDR tracks the target level within every species, and transfer is close to the identity among the animal species, conservative when calibrated on yeast or *E. coli* and applied to an animal species, and anti-conservative in the opposite direction.

**Fig. A1:**
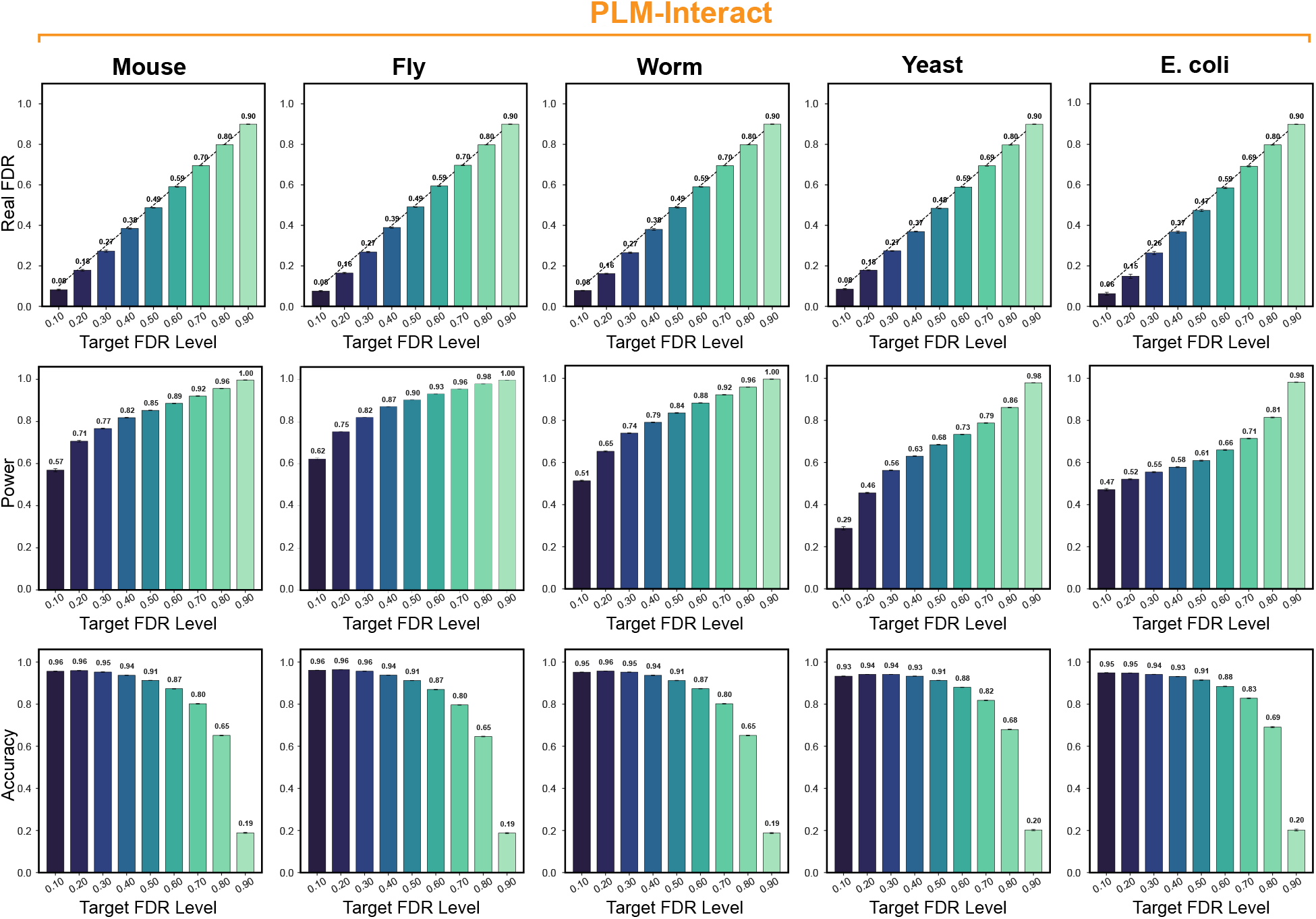
In-distribution calibration controls FDR across datasets with varying discovery power. Results are shown for PLM-Interact humanV12 across five datasets. Rows report empirical FDR, power, and accuracy over target FDR levels *q*. Bars show means over 100 random 50%/50% calibration and test splits with 95% confidence intervals. The dashed line marks empirical FDR equal to the target. Power is the fraction of true interactions in the test set that are selected.

**Fig. A2:**
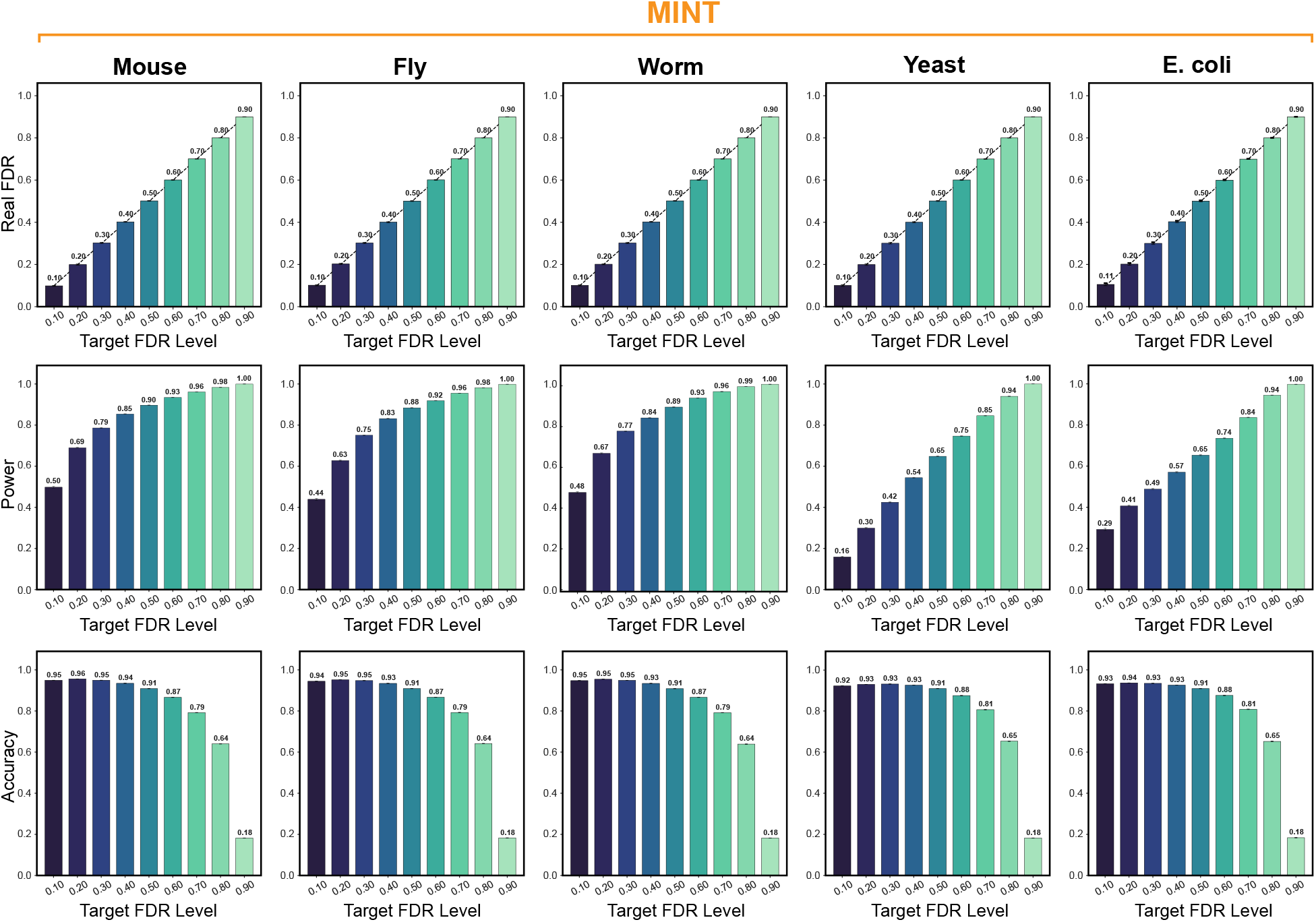
In-distribution calibration controls FDR across datasets with varying discovery power. Results are shown for MINT fine-tuned on STRING V11 human PPIs across five datasets. Rows report empirical FDR, power, and accuracy over target FDR levels *q*. Bars show means over 100 random 50%/50% calibration and test splits with 95% confidence intervals. The dashed line marks empirical FDR equal to the target. Power is the fraction of true interactions in the test set that are selected.

**Fig. A3:**
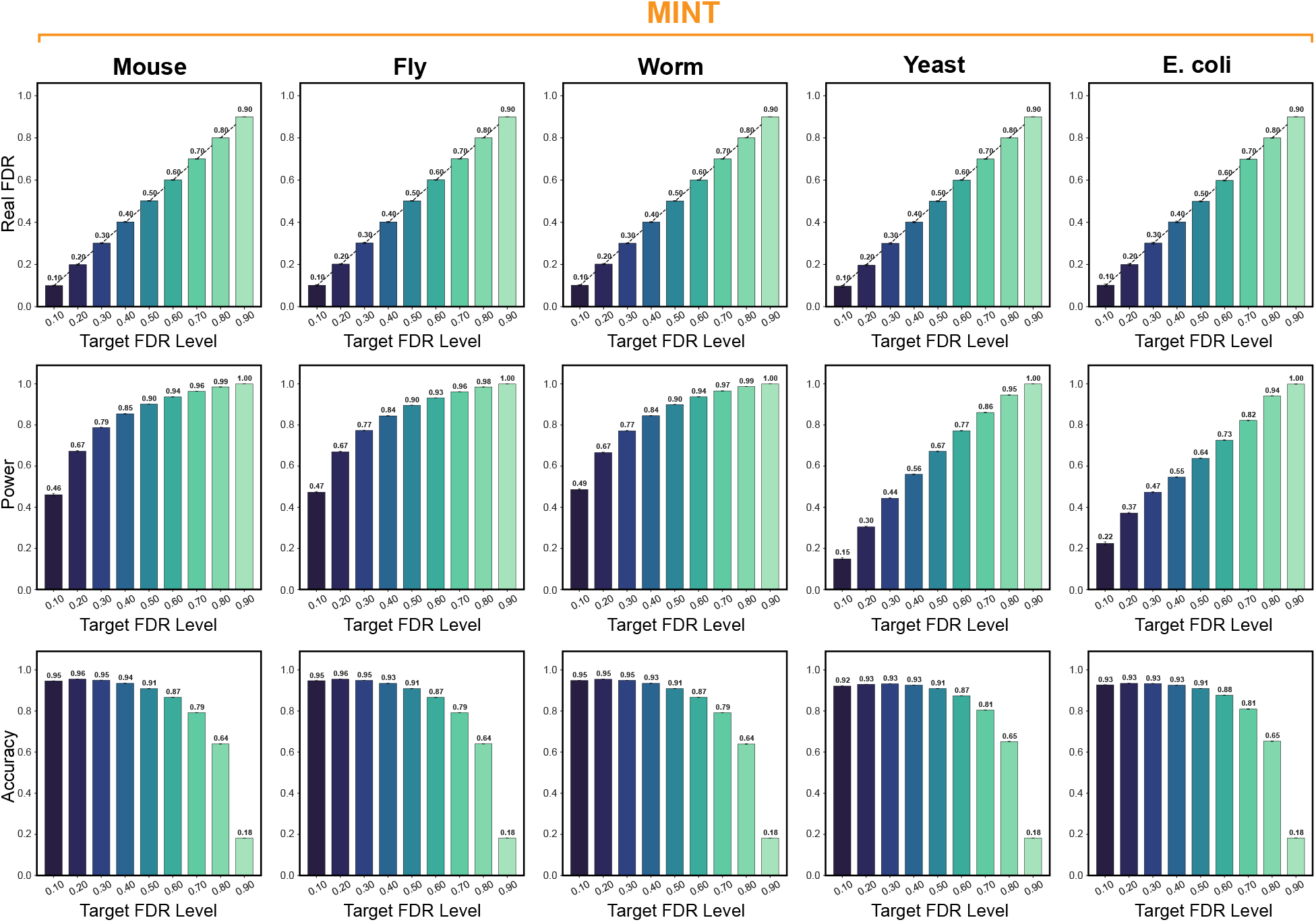
In-distribution calibration controls FDR across datasets with varying discovery power. Results are shown for MINT fine-tuned on STRING V12 human PPIs across five datasets. Rows report empirical FDR, power, and accuracy over target FDR levels *q*. Bars show means over 100 random 50%/50% calibration and test splits with 95% confidence intervals. The dashed line marks empirical FDR equal to the target. Power is the fraction of true interactions in the test set that are selected.

**Fig. A4:**
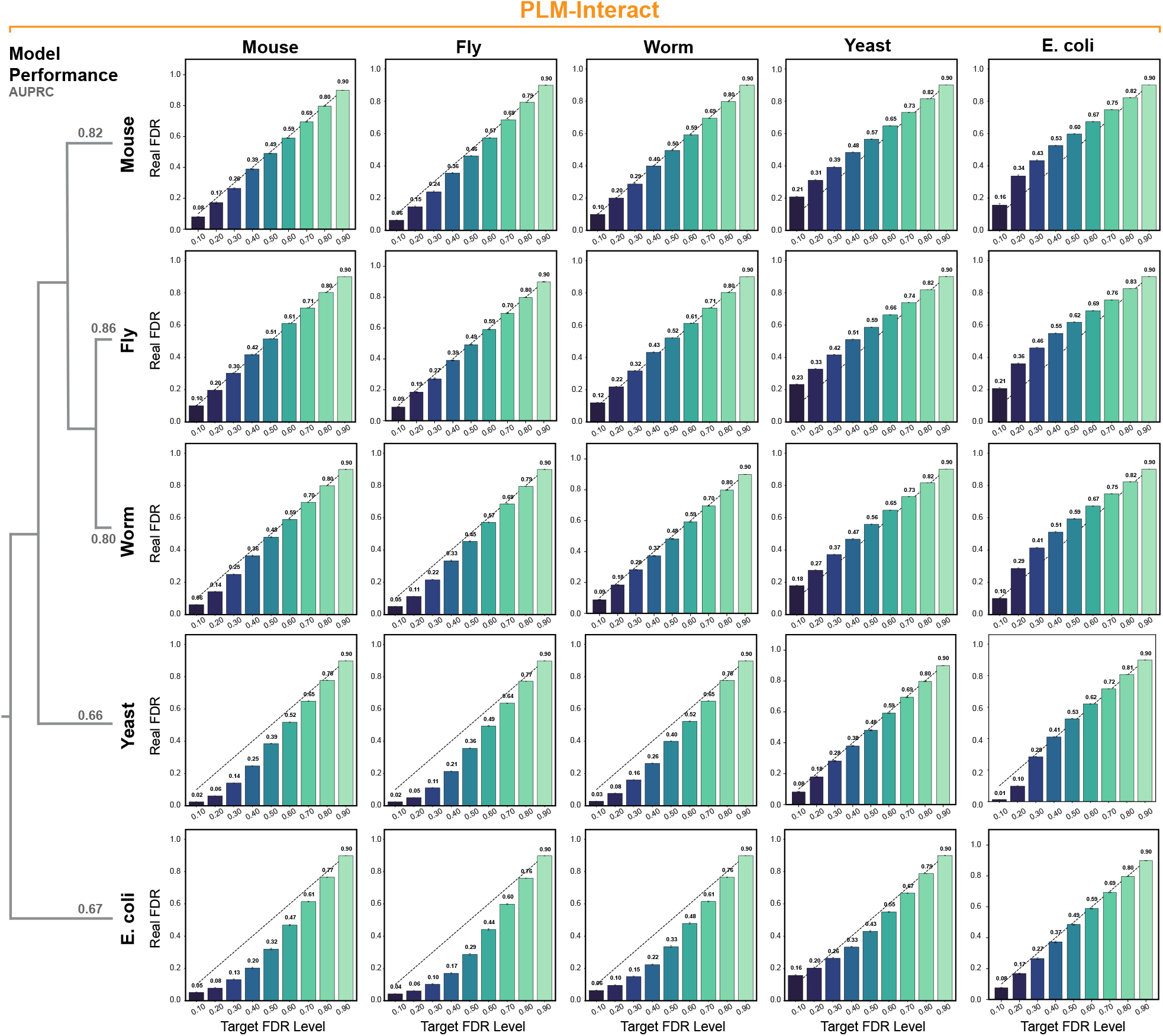
FDR control under distribution shift depends on transfer direction. Results are shown for PLM-Interact humanV12. Rows indicate calibration datasets and columns indicate test datasets, with diagonal panels showing in-distribution evaluation. Bars show mean FDP over 100 repetitions with 95% confidence intervals, and the dashed line marks the target FDR. AUPRC values summarize prediction performance on each dataset.

**Fig. A5:**
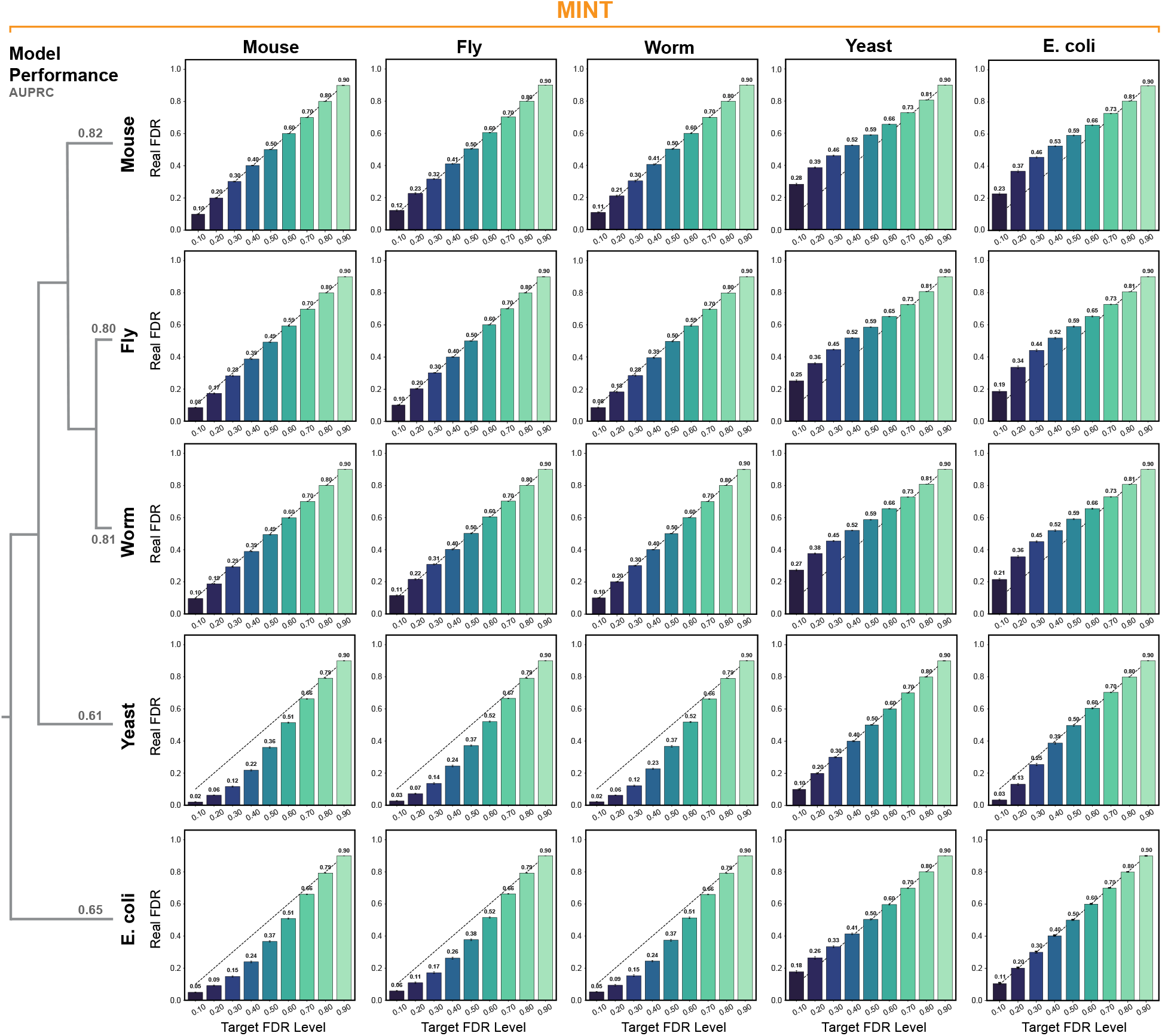
FDR control under distribution shift depends on transfer direction. Results are shown for MINT fine-tuned on STRING V11 human PPIs. Rows indicate calibration datasets and columns indicate test datasets, with diagonal panels showing in-distribution evaluation. Bars show mean FDP over 100 repetitions with 95% confidence intervals, and the dashed line marks the target FDR. AUPRC values summarize prediction performance on each dataset.

**Fig. A6:**
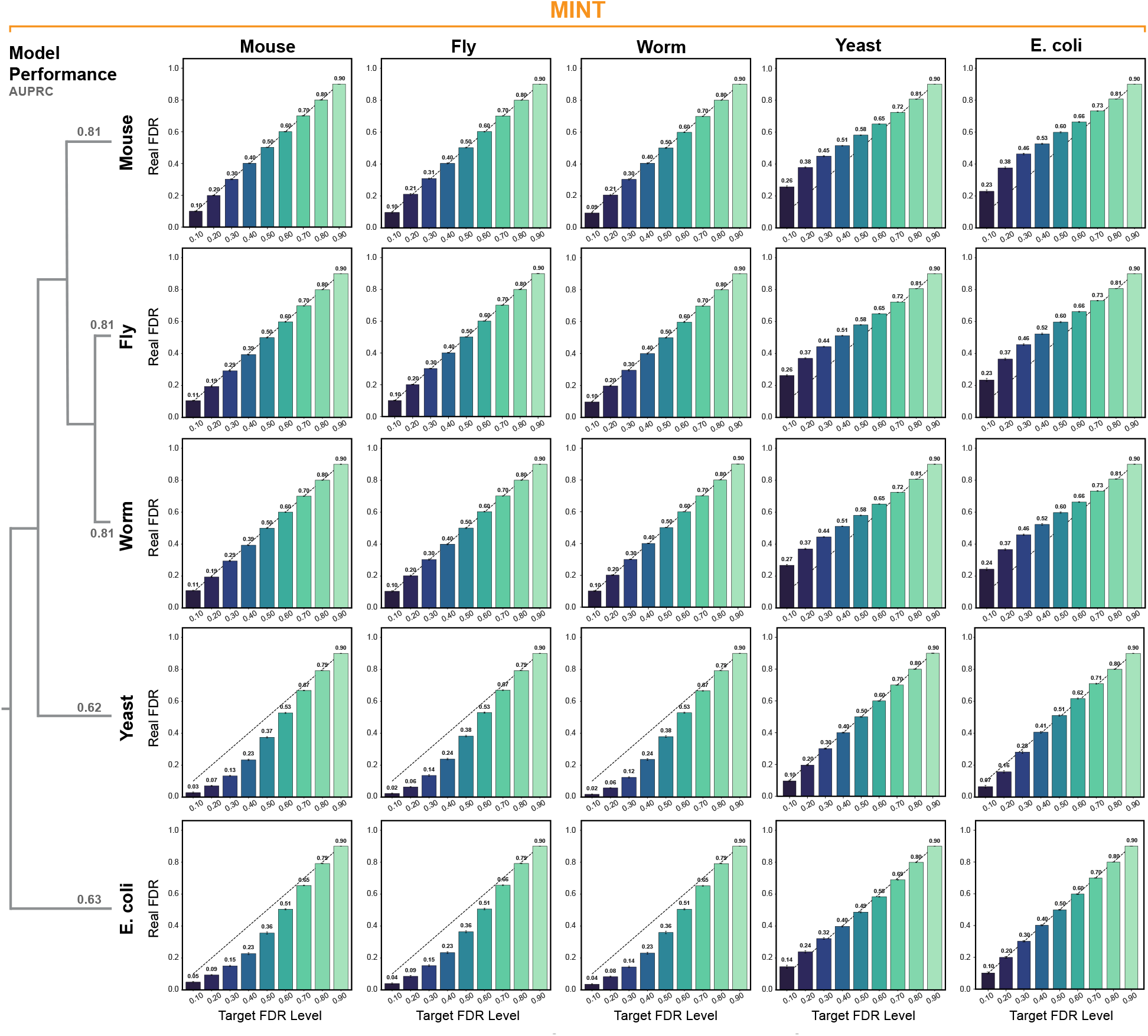
FDR control under distribution shift depends on transfer direction. Results are shown for MINT fine-tuned on STRING V12 human PPIs. Rows indicate calibration datasets and columns indicate test datasets, with diagonal panels showing in-distribution evaluation. Bars show mean FDP over 100 repetitions with 95% confidence intervals, and the dashed line marks the target FDR. AUPRC values summarize prediction performance on each dataset.

